# Actin Painting: a multicolor fluorescence staining method highlighting cell type-specific differences in the composition of filamentous actin architectures

**DOI:** 10.1101/2025.02.05.636544

**Authors:** Akira Nagasaki, Saku T. Kijima, Yoichi Shinkai, Yukio Ohtsuka, Tomoyo Ochiishi, Yasnory T.F. Sasaki, Saori Hata, Kazumi Hirano, Yoshio Kato, Motomichi Doi, Taro Q.P. Uyeda

## Abstract

Actin is a major structural component of the cytoskeleton in eukaryotic cells, and filamentous actin (F-actin) forms a variety of types of cellular structures. Many actin probes have been developed to visualize F-actin architectures in eukaryotic cells. However, it is known that double-stained images of F-actin obtained by two different types of actin probes often show partial inconsistencies. While developing actin probes, we observed that the merged images of three distinct actin probes, each labeled with a different fluorescent dye (red, green and blue), enabled various F-actin architectures to be distinguished based on color. This differentiation arises from slight variations in the distribution of each actin probe, and results in a unique color combination of actin probes that reflects the relative intensities of the three fluorescent dyes. In this report, we introduce a new cell staining method, named Actin Painting, which exploits the unique affinity of actin-binding molecules in each actin probe to F-actin architectures. This technique allows the classification of various cell types based on their specific actin cytoskeleton.

## Introduction

Actin is a highly ubiquitous proteins in eukaryotes, and monomeric actin proteins undergo polymerization to form microfilaments, filamentous actin (F-actin) within cells. F-actin gives rise to a wide variety of actin-based architectures, collectively referred to as the actin cytoskeleton, which encompasses structures such as stress fibers, contractile rings, a cortical actin meshwork, lamellipodia, filopodia, focal adhesion, and more. The actin cytoskeleton plays a critical role in various cellular events, including cell migration, adhesion, cell division, membrane traffic, and cell signaling. Moreover, the type, distribution, and ratio of F-actin architectures in a cell line are specific to the morphology and function of tissues and organs (Pollard & Cooper, 2009; Blanchoin *et al*, 2014; Rottner *et al*, 2017; Lim & Plachta, 2021).

There have been ongoing efforts to visualize intracellular F-actin since the 1970s, resulting in the development of various staining methods. The first attempt to stain the actin cytoskeleton was performed by an immunofluorescent method with a specific anti-actin antibody in non-muscle cells (Lazarides & Weber, 1974). A separate approach involved the development of actin-binding molecules with a fluorescent dye that were used to detect of actin filaments, as described next. For example, fluorescein-labeled S1 fragments of myosin II allowed actin filaments to be visualized in cultured cells (Schloss *et al*, 1977). In 1979, a staining method that employed a derivative of phalloidin labeled with fluorescent dyes was developed (Wulf *et al*, 1979). Since then, this method has been widely adopted as the gold standard method for F-actin staining.

The advent of GFP technology in the late 1990s enable the real-time observation of the dynamic and distribution of proteins in living cells. EGFP-actin fusion protein was developed at an early stage of the development of GFP technology, and has been widely used for F-actin staining (Choidas *et al*, 1998). Subsequently, several probes to visualize the actin cytoskeleton were developed by fusing fluorescent protein and various types of actin-binding domains (ABD) (Melak *et al*, 2017). Among these, Lifeact-EGFP was frequently used to observe F-actin in eukaryotic cells (Riedl *et al*, 2010). The actin-binding motif, Lifeact, is a 17 amino acid peptide derived from an actin-binding protein, Abp140, of *Saccharomyces cerevisiae*, and is the smallest actin probe among the GFP type of fusion proteins.

As was mentioned above, various staining methods for F-actin have been developed, and these actin probes can be categorized into two types (Melak *et al*., 2017). The first type (type I) includes probes in which fluorescent dyes directly bind to actin protein, while the second type (type II) includes probes in which fluorescent dyes bind to actin-binding molecules. Type II actin probes are designed by combining actin-binding molecules with fluorescent dyes. It is generally recognized that actin probes show the same distribution of F-actin in fixed or living cells (Vazquez-Victorio *et al*, 2016).

However, slight differences in staining images by each Type II actin probe have been noted (Washington & Knecht, 2008; Uyeda *et al*, 2011; Belin *et al*, 2014; Sliogeryte *et al*, 2016; DesMarais *et al*, 2019; Harris *et al*, 2020; Mazloom-Farsibaf *et al*, 2021). These observation-driven findings are not inconsistent with the notion that each actin-binding molecule has a unique affinity to each F-actin architecture. Hence, focusing on these binding specificities among actin probes, in this study, we utilized the subtle differences in staining patterns of each actin probe across various types of each F-actin architecture to develop a novel molecular and cellular approach. We name this new method Actin Painting, which is based on additive mixing of the three primary colors, as triple-merged images of three different actin probes with different fluorescent dyes. This method enables the identification of distinct types of F-actin architectures based on their color.

## Material and Methods

### Cell culture

U2OS, COS-7, HeLa, HEK293T, MCF-7, and HT-1080 cells were cultured in Dulbecco’s Modified Eagle Medium (Fujifilm, Osaka, Japan) containing 10% fetal bovine serum (Daiichi Pure Chemicals, Tokyo, Japan) and penicillin streptomycin solution (Fujifilm). NBT-L2b, NBT-T1, and MDA-MB-231 were cultured in Eagle’s Minimum Medium (Fujifilm) containing 10% fetal bovine serum, a non-essential amino acid mixture (Fujifilm), and penicillin streptomycin solution (Fujifilm).

### Preparation of actin probes

Alexa Fluor Plus 405 phalloidin, fluorescein phalloidin, rhodamine phalloidin and phalloidin-iFluor 405 were purchased from Thermo Fisher Scientific (Tokyo, Japan), Cytoskeleton Inc (Denver, CO, USA), Fujifilm and Abcam (Tokyo, Japan), respectively. For the expression of recombinant proteins, a cold-shock expression vector pColdI (Takara Bio, Shiga, Japan) was used. PCR fragments of each ABD and fluorescent protein cDNA were cloned into the multi-cloning sites of pColdI. The pColdI vector expressing fusion protein of the ABD and fluorescent protein were transformed into competent Rosetta (DE3) *Escherichia coli* cells (Novagen, Tokyo, Japan). *E. coli* cells were cultured in 2xYT medium with 100 μg/mL ampicillin at 37°C and an optical density of 0.5 at 600 nm. The expression of recombinant proteins was induced by adding 1 mM IPTG and culturing *E. coli* cells at 16°C for 15 h. Recombinant proteins were purified on Ni-NTA agarose (Qiagen, Tokyo, Japan). The list of the actin probes used as well as their amino acid sequences, are shown in supplementary Table 1.

### Actin Painting for cultured cells and primary cultured neurons

Established cell lines were cultured on collagen coated glass coverslips in each specific medium for 24 h. Primary neurons from mouse embryos were cultured on a poly-L-lysine-coated coverslip (Matsunami, Tokyo, Japan) in Neurobasal medium (Life Technologies, Tokyo, Japan) for two weeks. Cells were fixed with fixation buffer, 1x PBS (10x D-PBS (-), Fujifilm) containing 0.5% formaldehyde and 1mM MgCl_2_ for 15 min at room temperature. After washing three times with wash buffer (PBS containing 1mM DTT and 1 mM MgCl_2_), cells were treated with a permeabilization buffer (PBS containing 0.1% Triton X-100, 1 mM DTT, and 1 mM MgCl_2_) for 5 min at room temperature. After washing three times with the wash buffer, cells were treated with 500 μL of staining solution containing one unit of phalloidin derivatives and 25-100 ug of fusion proteins in wash buffer for 60 min at room temperature. After washing three times with the wash buffer, cells were fixed again by PBS containing 6% formaldehyde and 1 mM MgCl_2_ for 15 min. Fixed cells were washed with the washing buffer, and the coverslip was transferred to a slide glass with a mounting medium containing 1.25% of 1,4 diazabicyclo [2,2,2] octane (Fujifilm), 2.5% of polyvinyl alcohol (average MW 1,500∼1,800, Fujifilm), 5% of glycerol (Fujifilm) and 50 mM Tris-HCl (pH8.5).

### Actin Painting for tissue sections

Frozen section slides of paraformaldehyde-fixed tissues were treated with the permeabilization buffer for 15∼30 min at room temperature. Tissue slides were then washed three times with wash buffer. For F-actin staining, actin probes were diluted in PBS buffer containing 1mM DTT, 1 mM, 1mM MgCl_2_, 1 mM ATP, and 1x protease inhibitor cocktail (Fujifilm). A mixture containing one unit of phalloidin derivatives and 25-100 ug of fusion proteins was added to tissue slides and incubated at 4°C for 12 h. After washing three times with PBS containing 1mM DTT, and 1 mM MgCl_2_, tissue slides were re-fixed with 8% formalin neutral buffer solution (Fujifilm) for 30 min. The fixation buffer was washed out and the slides were prepared using mounted with mounting medium by adding DMSO solution of protease inhibitor mixture as described above.

### Actin Painting for sea squirt larva, pharyngeal muscle of a nematode, and organoid

For actin staining of a sea squirt larva (*Halocynthia roretzi*), pharyngeal muscle of a freeze-cracking nematode (*Chaenorhabditis elegans*), and mature brain organoids, samples were fixed in 0.5% formaldehyde in PBS buffer containing 1mM MgCl_2_, for more than 3 h at 4°C, then incubated in the permeabilization buffer for 1 h at 4°C. After washing three times with washing buffer, the mixture containing one unit of phalloidin derivatives and 25-100 ug of fusion proteins was added to the samples and incubated at 4°C for 12 hours. After washing three times with the wash buffer, samples were re-fixed with 8% formalin neutral buffer solution (Fujifilm) for 30 min. The stained samples were observed on a glass-bottomed dish in PBS containing 15% glycerol, and 1.25% of 1,4 diazabicyclo [2,2,2] octane as an anti-fading agent.

## Results

### A mismatch in intracellular distribution among actin probes

We have developed actin probes for use in *Dictyostelium* (Asano *et al*, 2004; Uyeda *et al*., 2011; Shibata *et al*, 2016), plants (Kijima *et al*, 2018), and mammalian cells (Nagasaki *et al*, 2017). During the evaluation of actin probes across various cell types, we observed that images of actin filaments stained with phalloidin derivatives often did not completely correspond with those from other actin probes, although each actin probe showed a characteristic F-actin pattern. As shown in Figure 1, the F-actin image stained with Lifeact-EGFP did not exactly correspond to that with rhodamine phalloidin. The upper panels show cells expressing Lifeact-EGFP that were stained with rhodamine phalloidin after fixation and permeabilization. On the other hand, the lower panels show cells stained with purified Lifeact-EGFP protein and rhodamine phalloidin after fixation and permeabilization. All images show a typical actin distribution, but both merged images revealed a partial mismatch in localization (Figure 1A and B, right panels). The differences in distribution with rhodamine phalloidin seems to be more noticeable in cells post-stained with Lifeact-EGFP, as demonstrated by line scan results, compared to cells transfected with Lifeact-EGFP (Supplementary Figure 1A and B).

**Figure 1.**
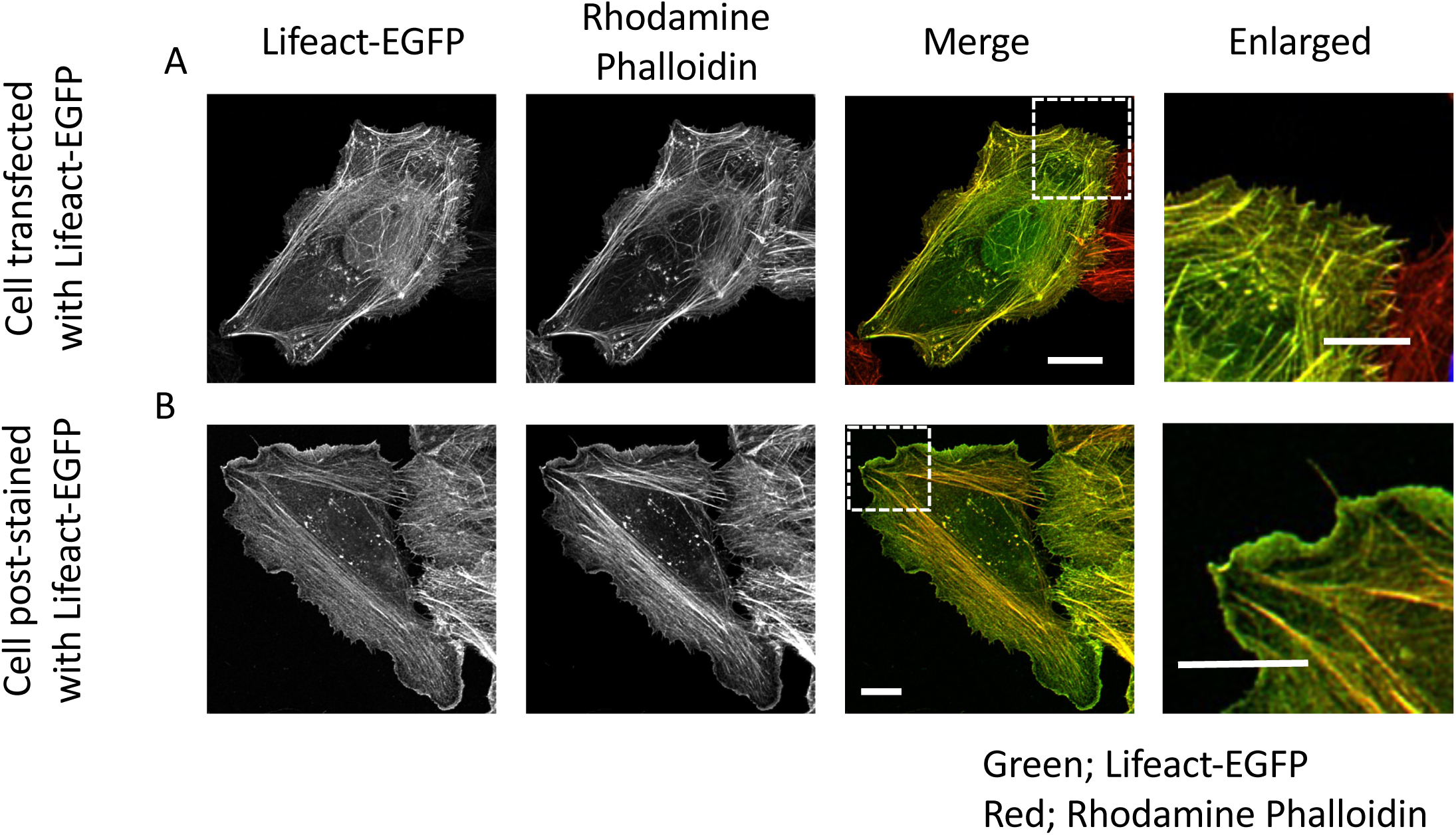
The merged images of double staining with Lifeact-EGFP and Phalloidin (A) A U2OS cell expressing Lifeact-EGFP was fixed with formaldehyde and treated with Triton X-100. Fixed transfected cells were stained with rhodamine-phalloidin. Non-transfected cells are stained red. (B) A fixed U2OS cell was stained with rhodamine-phalloidin and purified Lifeact-EGFP. Both images of Lifeact-EGFP (Green) and rhodamine-phalloidin (Red) were merged to create a single image. These images were captured by confocal microscopy, NIKON A1 and were generated by the Z-projection plugin of ImageJ software. White boxed areas are enlarged on the right. Bar; 10 μm

### Merged images of three types of actin probes with different color fluorescent dyes

In this report, we focused on the subtle differential distribution among probes based on various actin-binding molecules, such as phalloidin, Lifeact and several ABDs, as was described in the introduction. After adding an additional third actin probe with a different fluorescent dye, it was possible to distinguish each kind of F-actin architecture with a distinct color on triple-stained actin images. Figure 2 shows merged actin images composed of fluorescence images of Alexa Fluor 405-phalloidn, purified EGFP-mouse talin1 ABD (EGFP-mTln ABD), and purified Lifeact-mCherry in a fixed U2OS cell. Lifeact and phalloidin are the widely used actin probes, and mTln ABD is conventional actin probe for *Arabidopsis thaliana* (Kost *et al*, 1998). Each actin probe image displayed typical actin filaments (Figure 2B, gray-scale images), although the color image highlighted slight differences in distribution among actin probes when the three images were merged. The different colors of each F-actin architectures in the merged image were caused by the specific combination and ratio of each actin probe that bound to the F-actin architectures, as shown in Supplementary Figure 2. When only one type of actin probe bound to F-actin, the merged images show the respective color, as either red, blue or green (Supplementary Figure 2A and C). On the other hand, the merged images of F-actin stained with two kinds of actin probes indicated cyan, magenta or yellow depending on the combination of probes. In addition, the merged images of F-actin labeled with the three kinds of actin probes indicated in white when the brightness and contrast of each image was adjusted. The differences in color of each F-actin architecture in cells result from the varying combinations of binding actin probes for each actin architecture, in accordance with an additive color theory, as shown on a color wheel (Supplementary Figure 2B). We named this staining method, which uses three types of actin probes with different fluorescent dyes, as Actin Painting.

**Figure 2.**
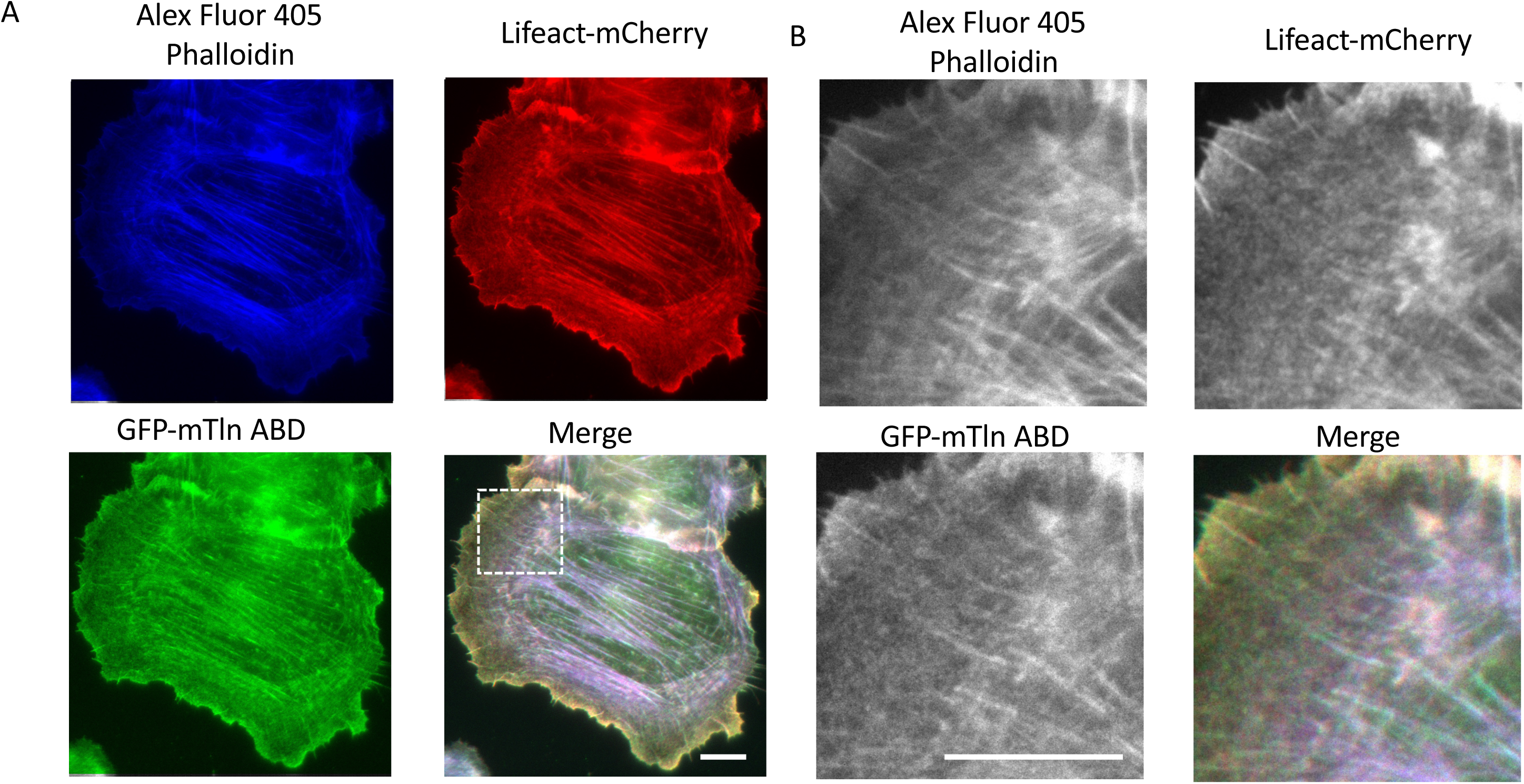
Triple staining of actin probes (A) A fixed U2OS cell was stained with Alexa Fluor 405 phalloidin (blue), purified GFP-mTln ABD (green) and purified Lifeact-mCherry (red). White boxed area is enlarged on the Panel (B). Images were captured by conventional fluorescent microscopy, Olympus IX81. Bar; 10 μm

The concentration of formaldehyde affected the effectiveness of staining by actin probes. Especially, the affinity of mTln ABD to F-actin was more sensitive to formalin fixation compared to phalloidin and Lifeact (Supplementary Figure 3A and B). In addition, type of fixation solution affected the pattern of staining of F-actin architectures (Supplementary Figure 4).

### Different combinations of actin probes for Actin Painting

We also investigated several combinations of actin probes for staining fixed U2OS cells. Since there are ample possible combinations of actin probes, we decided to focus on combinations of actin probes using phalloidin-iFluor 405 (blue), fusion protein of EGFP-ABD and fusion protein of mCherry-ABD (or cofilin). While each actin probe consistently displayed a typical F-actin pattern, the merged images highlighted differences in the affinities of probes to distinct F-actin architectures. The color patterns of F-actin architectures varied among cells stained with different probe combinations (Figure 3). The probe combinations, including GFP-mouse Talin1 ABD (mTln1 ABD), showed green focal adhesion at the tips of stress fibers (Figure 3A, B and E). Rng2 ABD, which is a calponin homology domain in the actin-binding protein of fission yeast, stains mainly thin stress fibers, together with phalloidin-iFluor 405 (Figure 3 C, D). Lifeact and its variant, Lifeact x2, a tandem repeat of 17 amino acids of the Lifeact peptide, exhibited different distribution patterns (Figure 3F). The distribution of Lifeact (E17K), a variant with improved binding to F-actin (Belyy *et al*, 2020), showed all F-actin architectures in cyan, merged blue and green, which largely corresponds to the distribution of phalloidin (Figure 3G, H and Supplementary Figure 5). The full length of cofilin protein can also be used as a probe in Actin Painting. Cofilin-mCherry protein predominantly stained the meshwork within lamellipodia, although cofilin protein binds to the surface of the coverslips and therefore tends to have a high background level (Figure 3E and J). These results indicated that the color tone can be adjusted by selecting different combinations of actin probes, depending on the purpose of staining. An indirect fluorescent antibody method that utilizes anti-actin antibodies also can also serve as a candidate actin probe for Actin Painting (Figure 3I). The amino acid sequences of the protein-based actin probes are shown in Supplementary Table 1.

**Figure 3.**
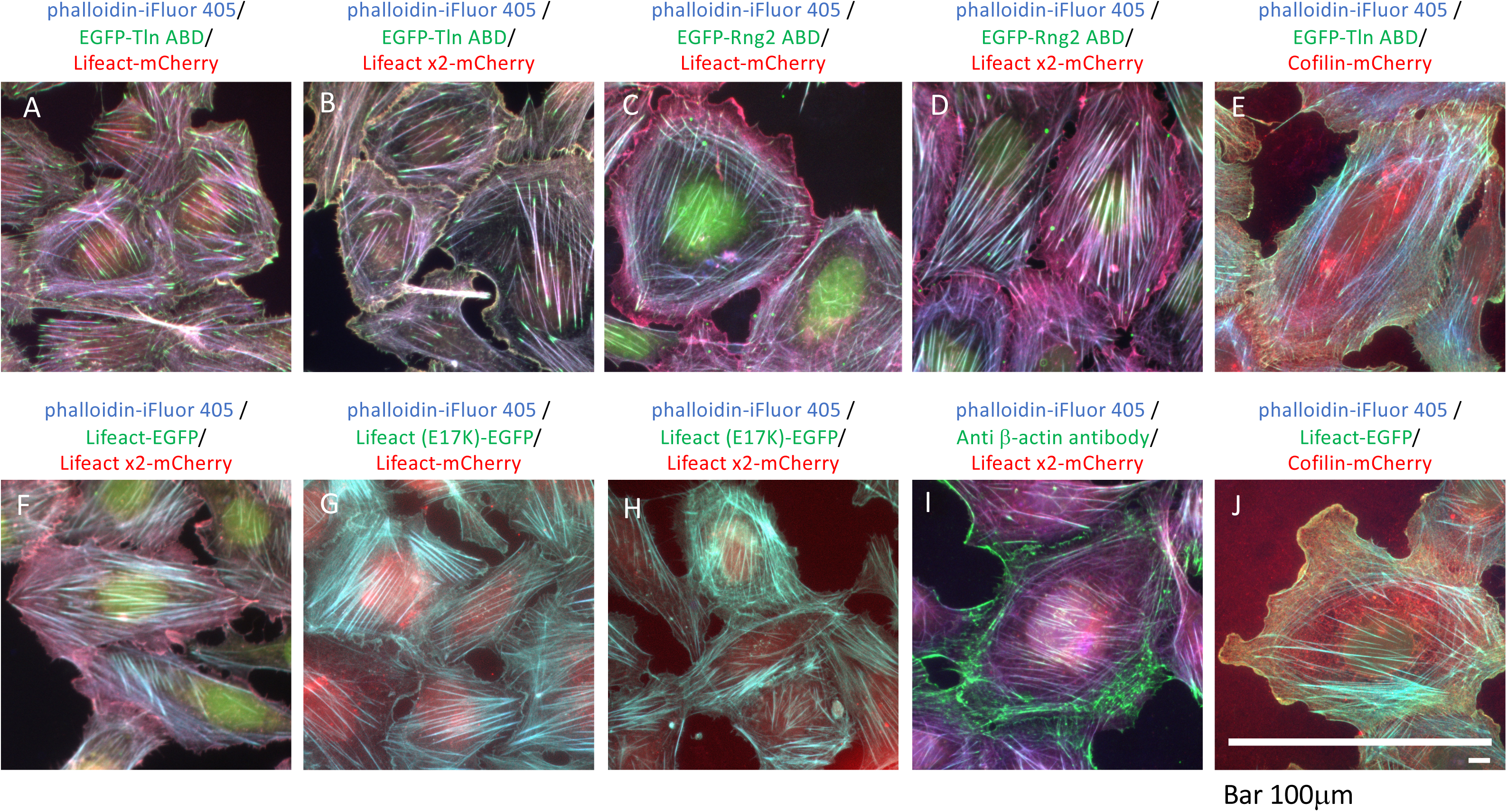
Staining with different combinations of actin probes Merged images of fixed U2OS cells stained with various combinations of three actin probes. All blue images were stained with phalloidin-iFluor 405 Reagents. Green and red show fluorescence of GFP and mCherry, respectively. Lifeact peptides are from *S. cerevisiae* (Riedl *et al*, 2008; Belyy *et al*., 2020). Rng2 ABD and mTln ABD are from *S. pombe* (Hayakawa *et al*, 2023) and *M. musculus* (Kost *et al*., 1998), respectively. Cofilin is from *D. discoideum* (Umeki *et al*, 2013). All images were captured by conventional fluorescent microscopy, Olympus IX81.Bar; 100 μm

### Staining of representative cell lines by Actin Painting

Several selected cell lines were stained by Actin Painting, and the actin probe combination used was phalloidin-iFluor 405, EGFP-mTln ABD, and Lifeact x2-mCherry. As shown in Figure 4, each cell line displayed a cell type-specific coloration using this combination of probes. These results suggest that the type and composition of F-actin architectures, including stress fiber, focal adhesion, filopodia, lamellipodia, *etc.*, are unique and specific to each cell line. For example, U2OS cells indicated green large focal adhesion and purple stress fibers. NBT-L2b had large lamellipodia including yellow F-actin, while HEK293 and MCF7 cells showed little diversity in F-actin architectures. In addition, when staining mouse primary neuronal cells by Actin Painting (Supplementary Figure 6), F-actin in the growth cone was stained in yellow while patch structures on dendrites were stained in green. This method allowed the growth cones and spines that formed in neuronal cells to be distinguish as yellow (white arrowheads) and green (white arrows), respectively. These results indicated that Actin Painting is able to detect differences in the composition of F-actin architectures in different cell lines based on differences in color.

**Figure 4.**
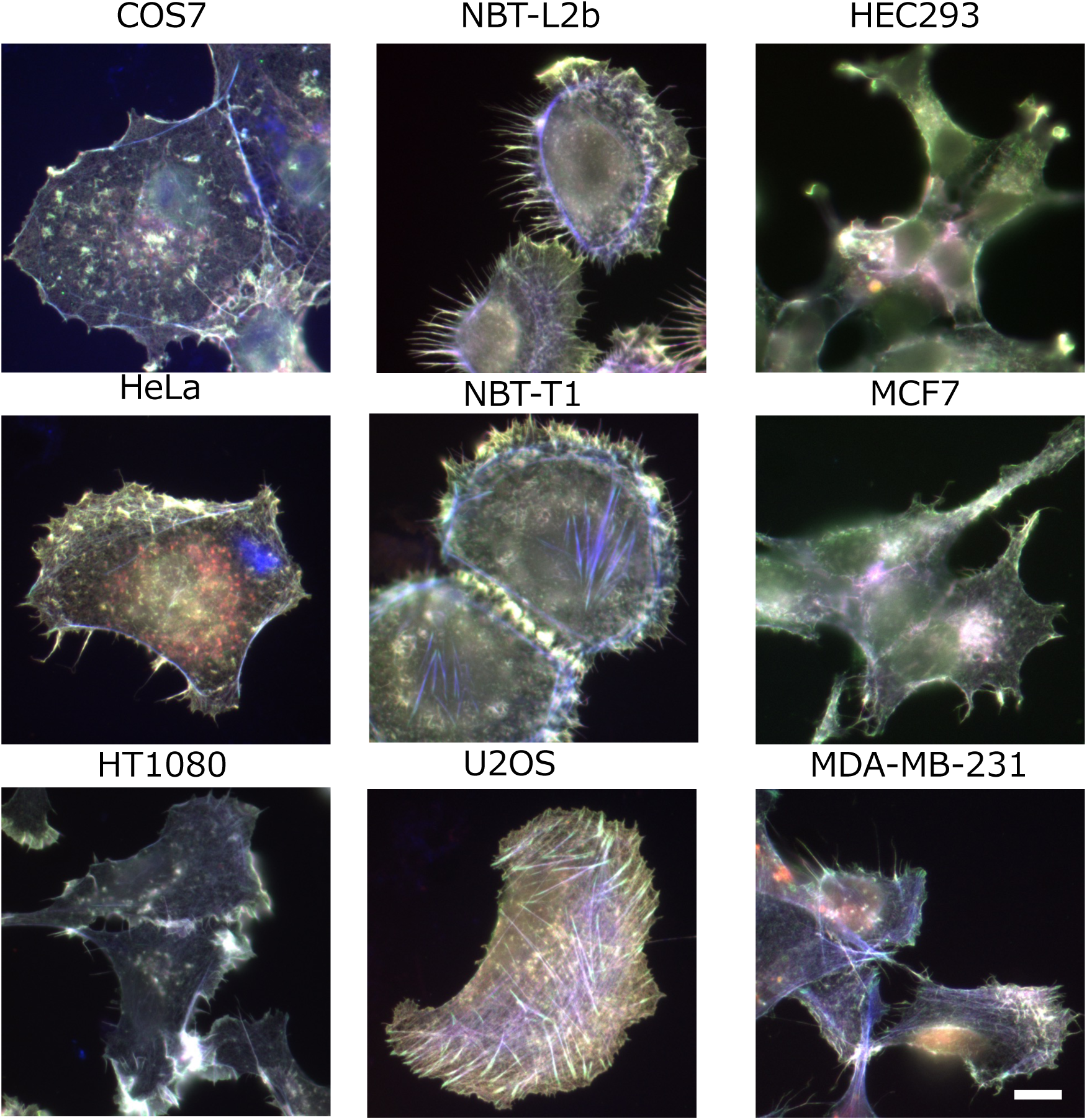
Several representative cell lines stained with Actin Painting Each cell was fixed with 0.5% formaldehyde and Triton X-100 on the coverslips. Cells were stained with phalloidin-iFluor 405 Reagents (blue), purified GFP-mTln ABD (green) and purified Lifeact x2-mCherry (red). Images were captured by conventional fluorescent microscopy, Olympus IX81. Bar; 10 μm

### Adaptation of Actin Painting for wound healing assay and co-culture

The wound healing assay is a general method for examining the directional migration of a cell population on substrates. Actin Painting was used to evaluate the wound healing assay. Figure 5A shows Actin Painting images of two cell lines, HeLa and MCF7, taken at 0 and 24 h after scratching. The gap formed by scratching with a pipette tip were filled with cells over time (Figure 5A, right). Actin Painting stained the cells along the edges facing the gap and other cells with distinct colors. These results indicated that the state of F-actin architectures was different in the cells along the edges and in those behind them. On the other hand, the yellow cells along the immediate sides of the gap at 0 h may have resulted from physical damage to the cytoskeleton caused by scratching.

**Figure 5.**
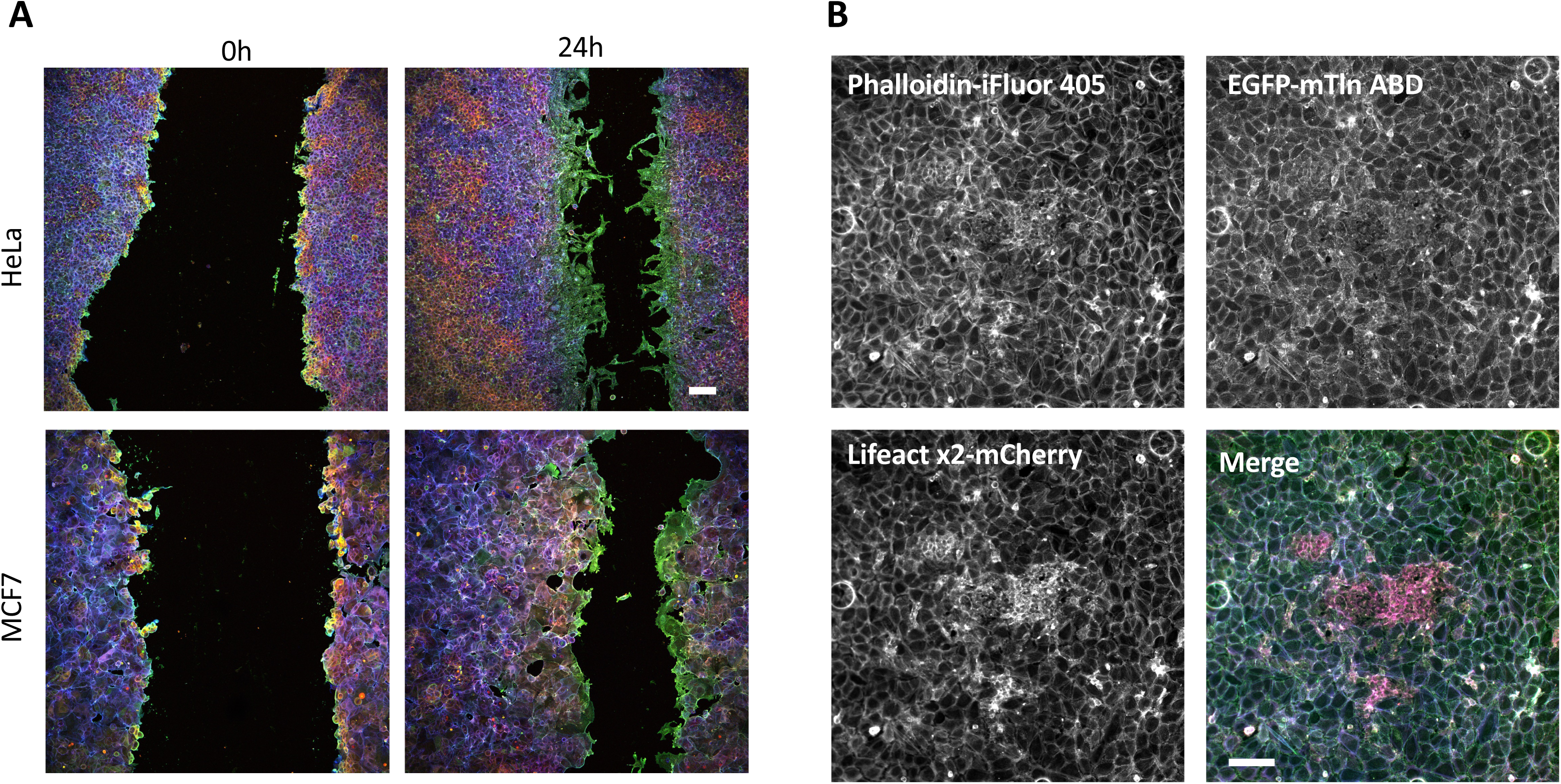
Actin Painting images of cells in wound healing assay and a co-culture of two different cell types (A) Merged images of wound healing assay of HeLa (upper row) and MCF7 cells (lower row). Fixed cells were stained with phalloidin-iFluor 405 Reagents (blue), purified GFP-mTln ABD (green) and purified Lifeact x2-mCherry (red) at 0 (left) and 24 hr (right) after scratching. Images were captured by confocal microscopy, NIKON A1. All images were generated by the Z-projection plugin of ImageJ software. Bar; 100 μm (B) Staining images of a co-culture of U2OS (blue-green) and HEK293 (magenta). Fixed cells were stained with phalloidin-iFluor 405 Reagents (blue), purified GFP-mTln ABD (green) and purified Lifeact x2-mCherry (red). Images were captured by conventional fluorescent microscopy, Olympus IX81. Bar; 100 μm

In addition, we attempted to stain a co-culture of U2OS cells and HEK293 cells that were cultured at a 10:1 ratio. Under these conditions, HEK293 cells formed colonies within the monolayer of U2OS cells on a glass-bottomed dish. Actin Painting highlighted the difference between cell types based on color: while U2OS cells in the monolayer appeared turquoise blue, colonies of HEK293 cells were magenta (Figure 5B). Compared to single probe staining, Actin Painting more clearly distinguished HEK293 and U2OS cells by distinct differences in color (Figure 5B, gray scale images and merged images).

### Staining of multicellular samples by Actin Painting

Tissues are composed of many different types of cells. It is known that many types of actin architectures based on filamentous actin formed in all cells of eukaryotes, leading to various cellular events depending on their respective functions. With this in mind, we attempted to stain multicellular tissues of a nematode, sea squirt, and organoids using Actin Painting. Each actin-binding molecule of the actin probes bound to several types of F-actin architectures according to their affinity to each of them. It was thus expected that various types of cells in each tissue would be stained by a distinct color using Actin Painting. The pharyngeal muscle of nematode, consisting of 95 cells across seven cell types, was stained in distinct colors for each cell type using this technique (Figure 6A). Actin Painting produced at least four different colors, red, blue, green and cyan. Figure 6B shows the Actin Painting-stained image of a sea squirt larva, indicating green, cyan, magenta and yellow from the head to the tail. The tail region of the sea squirt larva is composed of musculature and exhibits the characteristic patterns of sarcomeres (Figure 6B, white dotted square). Brain organoid was white with visually distinct round lumps (Figure 6C, right-side corner). However, Actin Painting revealed that brain organoids consisted of at least three cell types, as revealed by blue, green, and magenta staining (Figure 6C). These results indicated that Actin Painting can be adapted to detect different cell types in multicellular samples.

**Figure 6.**
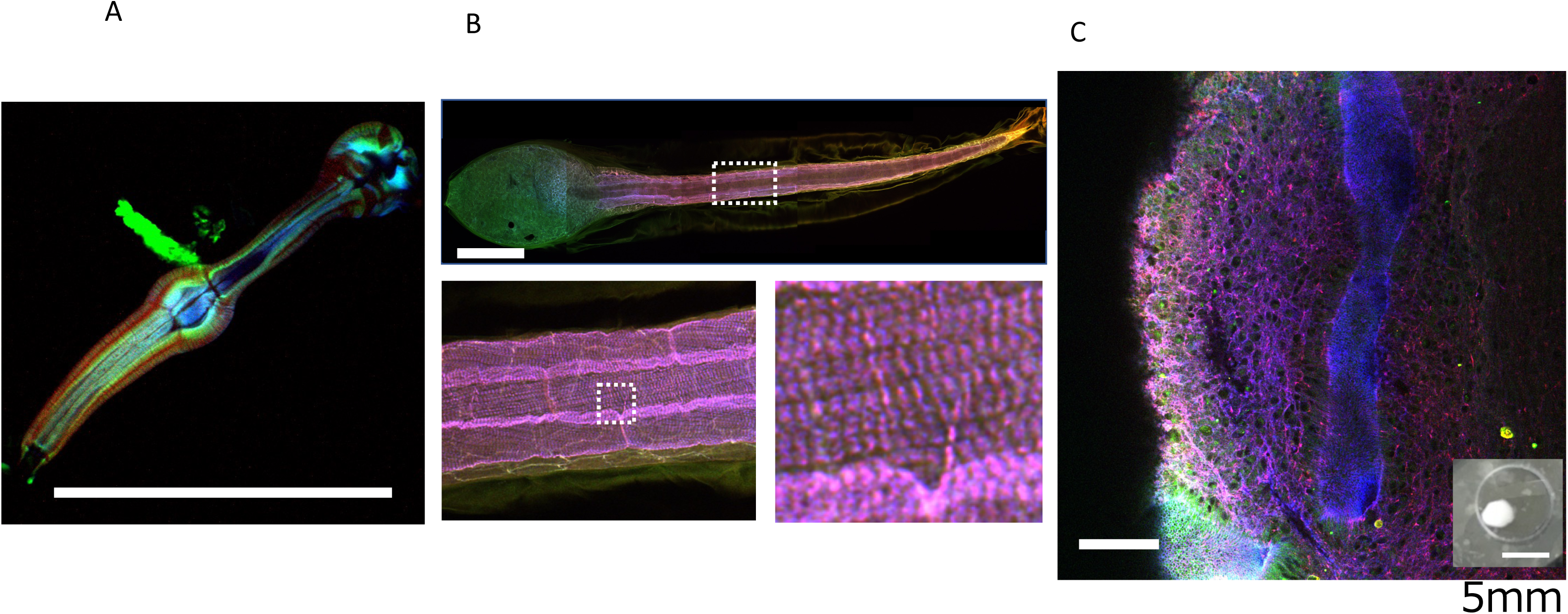
Merged images of multi-cellular samples stained with Actin Painting. Actin Painting was applied to (A) pharyngeal muscle of nematode, (B) larvae of sea squirt and (C) brain organoids. Fixed samples were stained with phalloidin-iFluor 405 Reagents (blue), purified GFP-mTln ABD (green) and purified Lifeact x2-mCherry (red). Actin Paint images of pharyngeal muscle of nematode was captured by Olympus conventional fluorescent microscopy IX81. Images of a larva of sea squirt and a brain organoid were obtained by Nikon confocal microscopy AI and each confocal images were converted into a single Z-projection plane with plugin of ImageJ software. Bar; 100 μm

### Staining of tissue slides by Actin Painting

Mouse tissue slides were stained with Actin Painting in an attempt to distinguish different cell types within the tissue based on distinct colors. Figure 7 shows Actin Painting images of a tissue slice of a mouse eyeball acquired by conventional fluorescence microscopy, with various cell types within the eye stained in distinct colors (Figure 7 center panel). The optic nerve stained green while the retina appears in various shades of green. Eye muscle and choroid are displayed in purple and cyan, respectively. The surrounding images are confocal Z-stack magnifications of specific regions within the central fluorescent image in Figure 7. Each confocal image revealed that various cells within the tissue formed a unique actin cytoskeleton, which were stained in different colors by Actin Painting, reflecting their specific functions. The original monochromatic images before merging are shown in Supplementary Figure 7. The variation in shading observed in each monochromatic image indicates differences in staining specificity of each probe for the distinct cell types in the mouse eye tissue.

**Figure 7.**
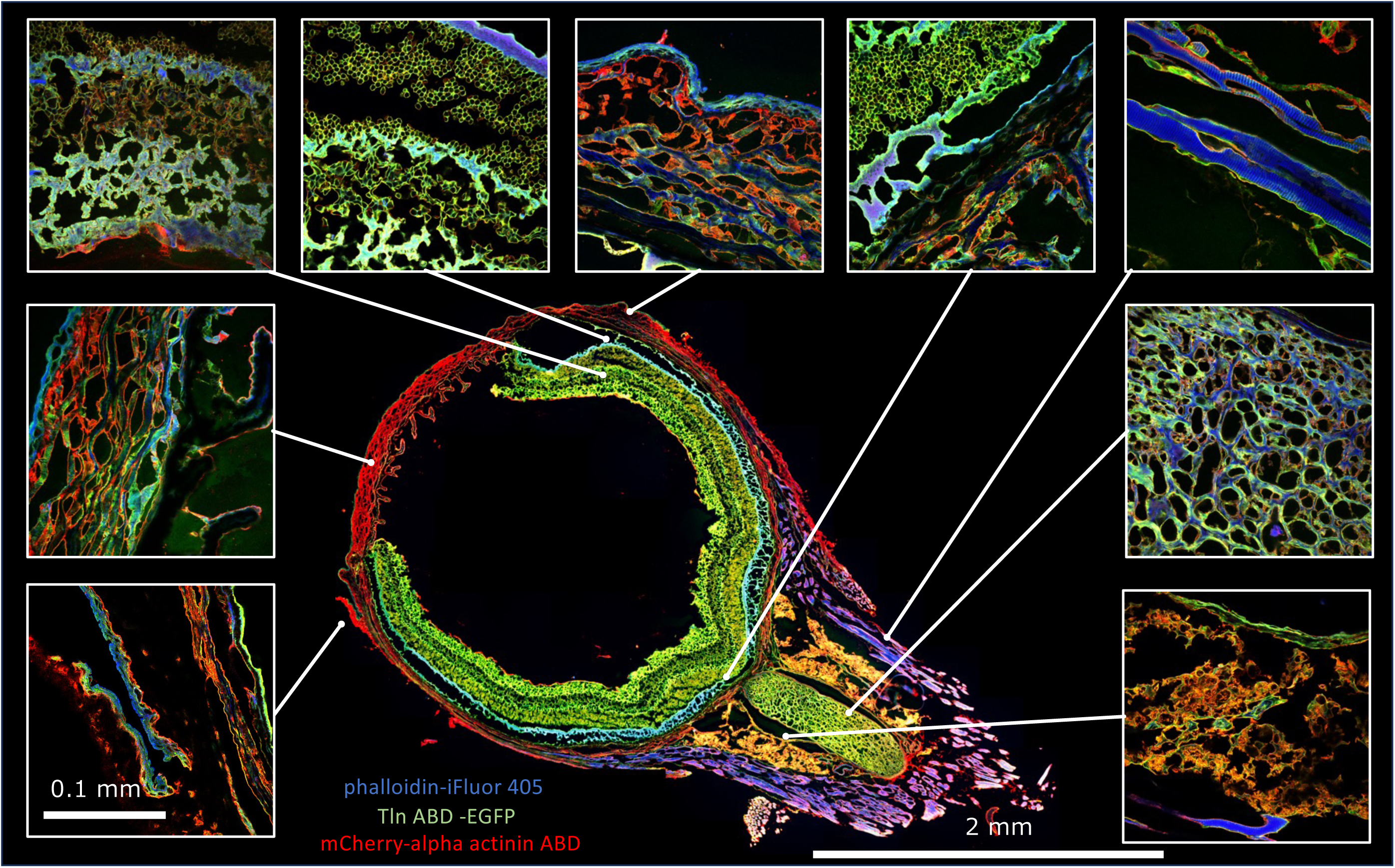
Actin Painting images of tissue slice of mouse eye. Tissue slice of a mouse eye was stained with phalloidin-iFluor 405 Reagents (blue), purified GFP-mTln ABD (Green) and purified *Dictyostelium* α-actinin ABD-mCherry (red). Images of mouse eye were captured by Olympus conventional fluorescent microscopy IX81, and 54 individual fluorescent images were stitched into a single composite image by Fiji software. Surrounding enlarged images were obtained by Nikon confocal microscopy AI. Scales are indicated on images.

## Discussion

Actin is essential for a variety of eukaryotic cellular processes, particularly in various cell motility mechanisms. Since the identification of cellular actin (Hatano and Oosawa, 1966), various actin staining techniques have been developed to visualize F-actin. For instance, immunofluorescence staining with specific antibodies for actin (Lazarides & Weber, 1974) and a phalloidin-labeling method (Wulf *et al*., 1979) have been widely applied for the visualization of F-actin in fixed cells. Over the years, fluorescent derivatives of phalloidin have become standard probes for cellular F-actin. In the late 1990s, the advent of GFP technology facilitated the detection of F-actin in living cells using various types of actin probes, including GFP-ABDs. Many detection methods for actin have been developed, and these can be categorized into two types. Type I actin probes involve fluorescent dyes that fuse directly to the actin protein, *e.g*., GFP-actin fusion protein (Choidas *et al*., 1998) or actin that is chemically labeled with a fluorescent dye (Taylor & Wang, 1978). On the other hand, Type II actin probes consist of fluorescent dyes fused to actin-binding molecules including phalloidin, Lifeact, antibodies, and various types of ABD from actin-binding proteins.

While all actin probes yield typical images of actin filaments, merged images resulting from double staining of different actin probes demonstrated that the localization of actin probes was not entirely consistent (Figure 1). A line scan of double-staining images of rhodamine phalloidin and Lifeact revealed similar patterns, although the heights of the respective peaks did not match (Supplementary Figure 1). These results indicated that each actin-binding molecule within an actin probe has unique binding abilities for distinct F-actin architecture. Specifically, it appears that rhodamine phalloidin preferentially binds to stress fibers in fixed U2OS cells relative to purified EGFP-Lifeact (Supplementary Figure 1). While difference in F-actin binding activities among actin probes have been recognized (Washington & Knecht, 2008; Uyeda *et al*., 2011; Belin *et al*., 2014; Sliogeryte *et al*., 2016; DesMarais *et al*., 2019; Harris *et al*., 2020; Harris *et al*, 2019; Mazloom-Farsibaf *et al*., 2021), phalloidin has generally been considered as an unbiased pan-actin probe. However, there are F-actin architectures that are resistant to staining by phalloidin. For example, it is known that phalloidin cannot bind to specific F-actin architectures, such as cofilin actin-rods (Munsie *et al*, 2009) or a certain type of nuclear actin filaments (Nagasaki *et al*, 2022). These results also demonstrate that each actin probe specifically recognizes distinct F-actin architectures with unique affinities, rather than providing uniform staining across all actin filaments.

As shown in Figure 2, it is unclear why a mismatched distribution among actin probes occurred for each F-actin architecture. However, the non-uniform staining among actin probes can be explained by the following five reasons: (1) actin isoforms, (2) post-translational modification, (3) binding of ABPs to F-actin, which may cause steric hindrance or structural changes, (4) different states of the ATPase domain in actin protein, and (5) structural distortion due to mechanical stress, including stretching and bending of actin protein. These factors result in actin protomers within each cellular F-actin architecture being structurally non-uniform, a phenomenon we refer to as cellular structural polymorphisms of F-actin. In mammalian cells, there are six actin isoforms with slightly different amino acid sequences at their N-termini. In addition, there are some reports that β-actin and γ-actin display isoform-specific F-actin architectures (Dugina *et al*, 2009; Simiczyjew *et al*, 2017) and distinct functions (Perrin & Ervasti, 2010; Pasquier *et al*, 2015). Furthermore, it is thought that the functional diversification of actin isoforms is achieved by isoform-specific post-translational modifications (Arora *et al*, 2023; Varland *et al*, 2019). One underlying causes of the mismatched distribution among F-actin probes may be due to their competitive interactions with ABDs, which are already bound specifically to actin filaments within their respective F-actin architectures. There are several examples, including filamin (Pavalko *et al*, 1989) in stress fibers, actinin 4 in focal adhesions (Kumeta *et al*, 2010) and Arps in lamellipodia (Johnston *et al*, 2008). On the other hand, cofilin binding to F-actin inhibits the binding of certain actin-binding molecules such as phalloidin and Lifeact (Munsie *et al*., 2009). In the case of cofilin, the binding sites of cofilin and phalloidin in F-actin are distinct, and phalloidin binding to F-actin is inhibited by structural alternation of phalloidin-binding sites that accompany the binding of cofilin to F-actin (McGough *et al*, 1997). Furthermore, Lifeact competes with cofilin and myosin II in a concentration-dependent manner (Belyy *et al*., 2020) and binding of the Arp2/3 complex to actin is inhibited by cofilin (Chan *et al*, 2009). Mechanical stress also affects the binding affinity of actin-binding molecules to F-actin. For example, we and others have reported that myosin II motor domains preferentially bind to mechanically stretched F-actin in *Dictyostelium* cells (Uyeda *et al*., 2011; Schiffhauer *et al*, 2016). Other mechanically regulated actin-biding proteins have been reported and summarized in a review (Sun & Alushin, 2023). It appears that cellular structural polymorphisms of F-actin cause the unique and non-uniform binding of respective actin probes to actin filaments in each F-actin architecture. The principle of Actin Painting is the visualization of different affinities among actin probes to different types of actin filaments by merging three kinds of actin probes. As shown in Supplementary Figure 2, the synthesized pseudo-color resulting from different combinations of actin probes binding to actin filaments in each F-actin architecture produced the Actin Painting images. Merged images of triple staining with phalloidin derivatives labeled with different fluorescent dyes displayed a white color by additive mixing of light (Supplementary Figure 8). This is because each phalloidin derivative has the same binding affinity to respective F-actin architectures.

Cell type-specific color patterns resulting from the use of Actin Painting can distinguish between two cell types in a co-culture of two cell lines (Figure 5B). Furthermore, the changes in actin cytoskeleton in a wound healing assay were visualized by a shift in color pattern (Figure 5A). This result indicates that Actin Painting is able to detect changes in the intracellular composition of F-actin architectures of individual cells, such as epithelial-mesenchymal transition, along a scratch. By staining with Actin Painting, individual cell lines show a unique color pattern, and these results indicated that each cell line has a unique combination of F-actin architectures that are specific to them. Actin Painting was then applied to multi-cellular samples, in the anticipation that it would be able to differentiate various cell types within organ tissues by producing distinct colors for each cell type. As expected, Actin Painting resulted in different colors for different cell types in a nematode pharynx, a sea squirt larva and an organoid (Figure 6). Actin Painting targets actin protein, which is present in large amounts in all eukaryotic cells. In comparison to similar existing techniques, such as multi-staining by immunofluorescent staining with antibodies for cell-specific markers, Actin Painting offers easy detection, ensuring that all cells are detectable, without exception.

Actin Painting can be adapted for staining tissue sections to generate multi-color images, as shown in Figure 7. A similar approach is multiplex immunohistochemistry staining, but multi-color staining with antibodies is generally limited to a display of seven colors due to constraints in the availability of fluorescent dyes and light sources, even when high-end fluorescence microscopy is used (Harms *et al*, 2023). In contrast, Actin Painting images can demonstrate a much wider range of colors by using a conventional fluorescence microscope equipped with a mercury lamp (Figure 7). Each of the three kinds of probes bound to various types of actin filaments in a specific proportion, resulting in a synthesis of colors by additive mixing of the three primary colors, red, green and blue, along with various intermediate colors (Figure 7). Moreover, the enlarged confocal image of Figure 7 reveals detailed F-actin architectures of each cell type in various colors, demonstrating that distinct F-actin characteristic in each cell type can be distinguished based on variations in color. Actin Painting demonstrates the diversity of the actin cytoskeleton, corresponding to a wide variety of cellular functions in respective cell types within organ tissues. In the future, simultaneous staining with antibodies against marker proteins will allow the identification of cell types using Actin Painting by correlating the observed color pattern with their respective cell type.

This method presents several challenges that need to be solved in the future. First, cellular staining by Actin Painting is influenced by the concentration of formaldehyde in fixation buffer (Supplementary figure 3). It is thought that a high concentration of formaldehyde inhibits the probe-specific binding of probes to F-actin due to masking of the binding sites. To ensure the specific binding of actin probes to F-actin, it is essential to avoid excessive cross-linking of amino acid residues of actin protein and excessive chemical modification of amino groups by aldehyde. Furthermore, the binding of actin probes to F-actin is affected by factors such as fixation condition, cell membrane permeabilization treatment, concentration of salt and pH of the buffer. Especially, fixation conditions are particularly crucial for Actin Painting. As demonstrated in Supplementary Figure 4, fixation with methanol appeared to inhibit the binding of the probes to F-actin. It is assumed that actin probes are unable to bind to denatured actin filaments because actin probes recognize the specific structure of an actin protomer within intact actin filaments. Furthermore, fixation with ALTFiX (glyoxal-based fixing solution, FALMA, Tokyo, Japan) eliminated the distinct binding pattern of phalloidin and mTln ABD to specific F-actin architectures, and abrogated the actin-binding affinity of Lifeact. Similarly, fixation with glutaraldehyde inhibited the binding of mTln ABD and Lifeact but did not affect phalloidin (Supplementary Figure 4). These findings suggest that certain fixation methods may disrupt the structural integrity of actin protomers within specific F-actin architectures, leading to the loss of specific binding of the actin probe to the F-actin architectures or the loss of binding properties. Conversely, all probes recognized actin filaments in acetone-treated cells, indicating that actin protein is resistant to acetone treatment (Supplementary Figure 4). In the purification process of actin from rabbit muscle, acetone is used in the first step, yet the purified actin possesses biochemical activity (Feuer *et al*, 1948). Given these known results, it is not surprising that actin was not denatured by the acetone treatment, allowing actin probes to bind to F-actin in cells treated with acetone.

Second, an attempt was made to apply Actin Painting to live cell imaging. However, it was challenging to observe the actin cytoskeleton in live cells expressing two types of actin probes. As shown in the lower panel of Supplementary Figure 9, the expression of Lifeact and mTln ABD led to the aggregation of actin probes in cells. We aim to identify the combination of actin probes that would allow F-actin to be monitored without aggregation in living cells. Recently, two actin probes, DARPin and Lifeact, were combined to simultaneously label F-actin architectures in living cells (Ivanova *et al*, 2024). Of interest, as shown in Supplementary Figure 9A, the expression of Lifeact-EGFP and Lifeact x2-mCherry in living cells eliminated the distribution mismatch observed in fixed cells post-stained with purified actin probes (Figure 3F). Post-staining with the purified actin probes may enable the detection of subtle differences in cellular structural polymorphisms of F-actin.

In addition, the fluorescence properties of lenses, detectors, optical filters and light sources influence color tones when RGB images are merged. It is also necessary to account for crosstalk in each channel, considering the spectrum of the fluorescent dye and the filter characteristics of the microscope. The development of standardized optical systems and reference samples for Actin Painting will be essential for enabling widely adopted quantitative analyses in the future.

In this report, we introduced new method named Actin Painting, which is based on staining of the actin cytoskeleton. This method is expected to be applicable in a variety of situations but staining and observation conditions influence the results of cellular staining. While we endeavored to optimize Actin Painting under limited conditions, we recognize that this method is still under development for general use. Further optimization will be necessary to expand Actin Painting for various quantitative assays in the future.

## Abbreviations

ABD: actin-binding domain
EGFP: enhanced green fluorescent protein
F-actin: filamentous actin

## Acknowledgements

This research was supported by the Management Expenses Grants from AIST and AMED under Grant Number JP23ym0126803 and the Japan Society for the Promotion of Science under Grant KAKENHI 24K22414. We are grateful to K. Katoh for technical support in microscopy work in this paper.

## Figure Legends

**Supplementary Figure 1. Dual staining of actin probes**

(A) Fixed U2OS cells were stained with purified Lifeact-EGFP and rhodamine phalloidin (Left). U2OS cells expressing EGFP-Lifeact were fixed and stained with purified Lifeact-EGFP and rhodamine phalloidin (Right). F-actin of non-transfected cells were stained red derived from rhodamine phalloidin in Right panel (white arrow). Nuclei were stained with Hoechst 33342 (blue). Images were captured by Olympus conventional fluorescent microscopy IX81. Bar; 10 μm (B) Line scan of cells stained with Lifeact-EGFP (green) and rhodamine phalloidin (red).

**Supplementary Figure 2. Scheme of the principle of Actin Painting**

(A) Combination of binding to an actin filament of three actin probes with different fluorescent dye. (B) Additive color wheel diagram. This diagram indicates that mixing two of the three primary colors, red, green, blue generates second color, cyan, magenta, yellow, and mixing of the three prime colors generates white. (C) Combination of actin probes binding to actin filaments and indicating color in the merged images. By combination of actin probes with different fluorescent dyes, actin filaments labeled with different ratios of the three probes exhibit different colors depending on their combinations.

**Supplementary Figure 3. Concentration of formaldehyde affected staining of Actin Painting**

(A) U2OS cells were fixed with formaldehyde at five different concentrations and permeabilized with 0.1% Triton X-100. Fixed cells were then stained with phalloidin-iFluor 405 Reagents, EGFP-mTln ABD, Lifeact x2-mCherry. Bar; 10 μm (B) Split channel images of U2OS cells fixed with 8% formaldehyde. Images were captured by Olympus conventional fluorescent microscopy IX81. Bar; 10 μm

**Supplementary Figure 4. Fixation methods affected staining of Actin Painting**

U2OS cells were fixed with 0.5% formaldehyde (FA), 100% methanol (MeOH), 1.0% glutaraldehyde (1.0% GA), Glyoxal-based, formalin substitute fixing solution (ALTFiX) and 100% acetone. Cells fixed with FA, GA and ALTFIX were permeabilized with 0.1% Triton X-100 to make the cell membrane permeable. Fixed cells were stained with phalloidin-iFluor 405, EGFP-mTln ABD, Lifeact x2-mCherry. All images were captured by Olympus conventional fluorescent microscopy IX81. Bar; 100 μm

**Supplementary Figure 5. Staining with Lifeact (E17K) and Phalloidin**

Fixed U2OS cells were stained with purified Lifeact (E17K)-EGFP and iFluor Phalloidin. The merged F-actin image is displayed in cyan due to the matching of each staining image. Images were captured by Olympus conventional fluorescent microscopy IX81.Bar; 100 μm

**Supplementary Figure 6. Actin Painting images of primary culture of mouse neurons**

Primary culture of mouse neurons on poly-L-lysine coated coverslips were fixed with formaldehyde and were permeable with 0.1% Triton X-100. Then, fixed cells were stained with phalloidin-iFluor 405 Reagents, EGFP-mTln ABD, Lifeact x2-mCherry. White boxed area in Left panel is enlarged. In right panel, growth cones and spines that formed in neuronal cells were stained yellow (white arrowheads) and green (white arrows). Images were captured by Olympus conventional fluorescent microscopy IX81. Scales are indicated on images.

**Supplementary Figure 7. Monochrome images of a tissue slice from the mouse eye stained with Actin Painting**.

The tissue slice was stained with iFluor 405-Phalloidin, mCherry-alpha actinin ABD and EGFP-mTln ABD. Merged images of them were indicated in Figure 7. Bar; 2 mm

**Supplementary Figure 8. Triple staining of phalloidin derivatives**

U2OS cells were stained with Alexa 405 phalloidin (A), Fluorescein Phalloidin (B,) and Rhodamine phalloidin (C). A merged image of triple staining (D). Images were captured by Olympus conventional fluorescent microscopy IX81. Bar; 100 μm

**Supplementary Figure 9. Living cells expressing two types of actin probes**

(A) U2OS cells expressing Lifeact-EGFP and Lifeact x2-mCherry. (B) U2OS cells expressing EGFP-mTln ABD and Lifeact x2-mCherry. Images were captured by Olympus conventional fluorescent microscopy IX81. Bar; 10 μm

**Supplementary Table 1. Information on protein-based actin probes**.

The table lists amino acid sequences of the proteins used in this report. Sequences are shown in the single letter codes.

