## Supplementary figures and images for "Actin Painting: a multicolor fluorescence staining method highlighting cell type-specific differences in the composition of filamentous actin architectures"

### Supplementary Figure 1

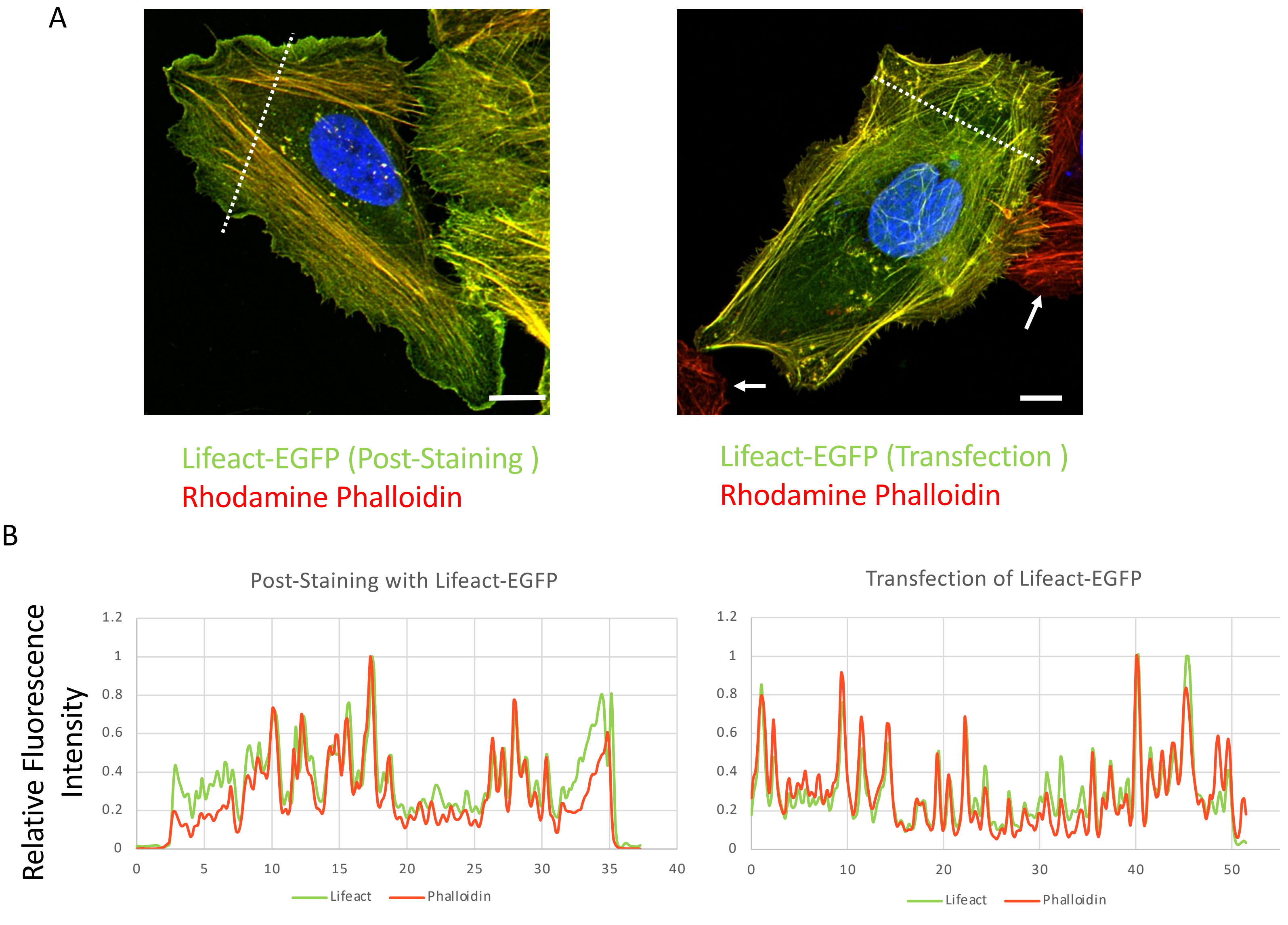


Nagasaki et al.,
Supp Figure 1

### Supplementary Figure 2

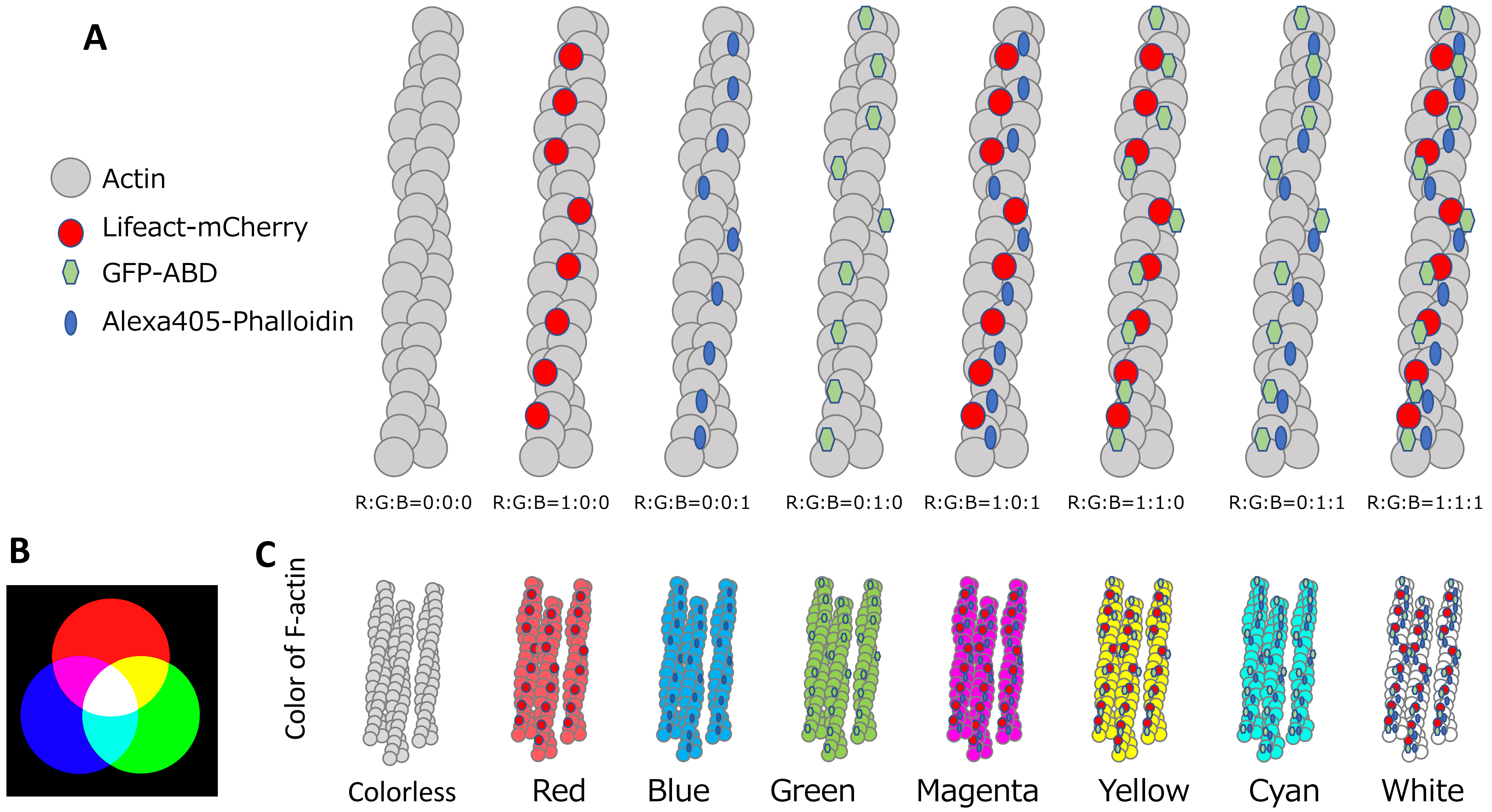


Nagasaki et al., Supp Figure 2

### Supplementary Figure 3

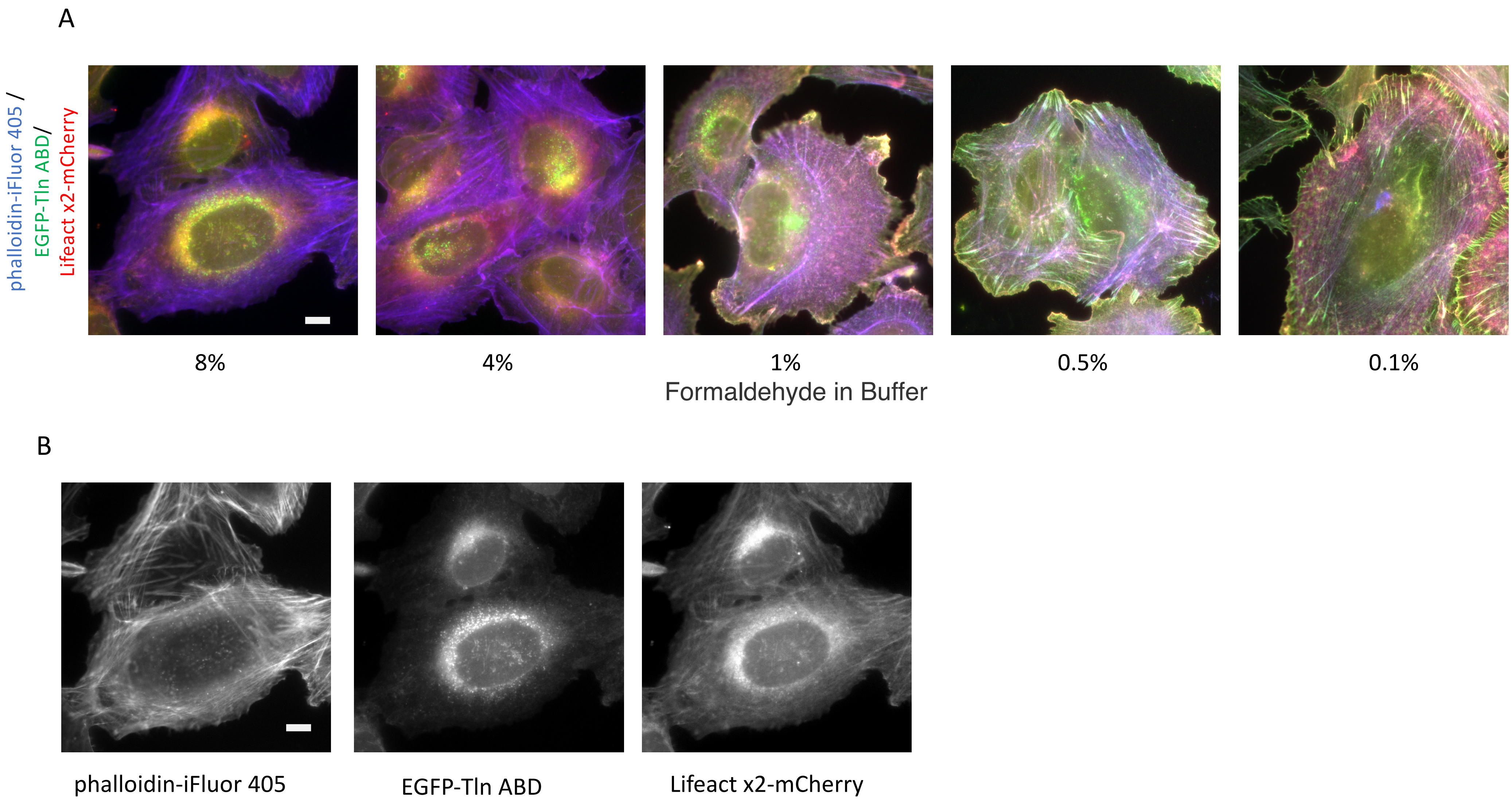


Nagasaki et al., Supp Figure 3

### Supplementary Figure 4

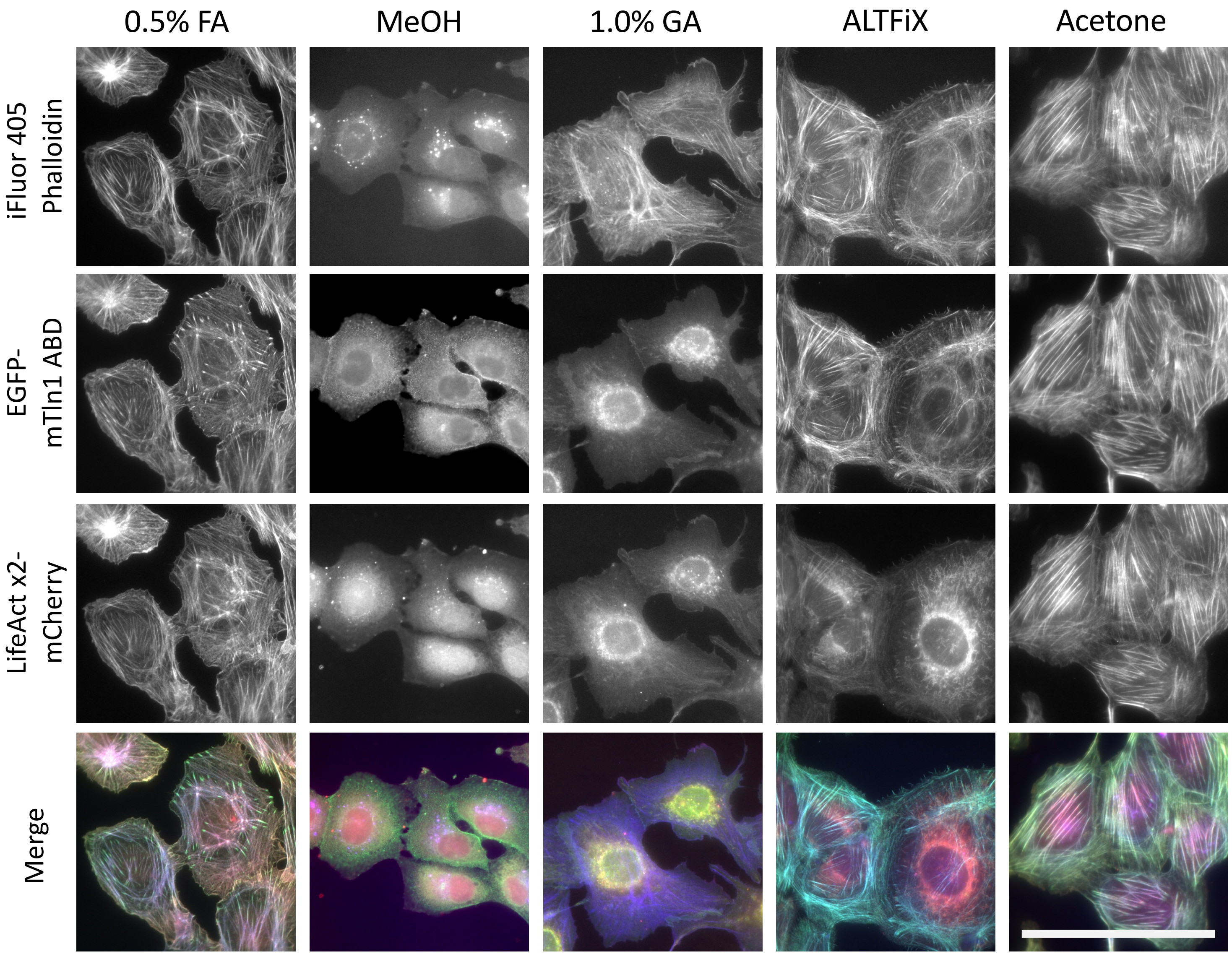


Nagasaki et al.,
Supp Figure 4

### Supplementary Figure 5

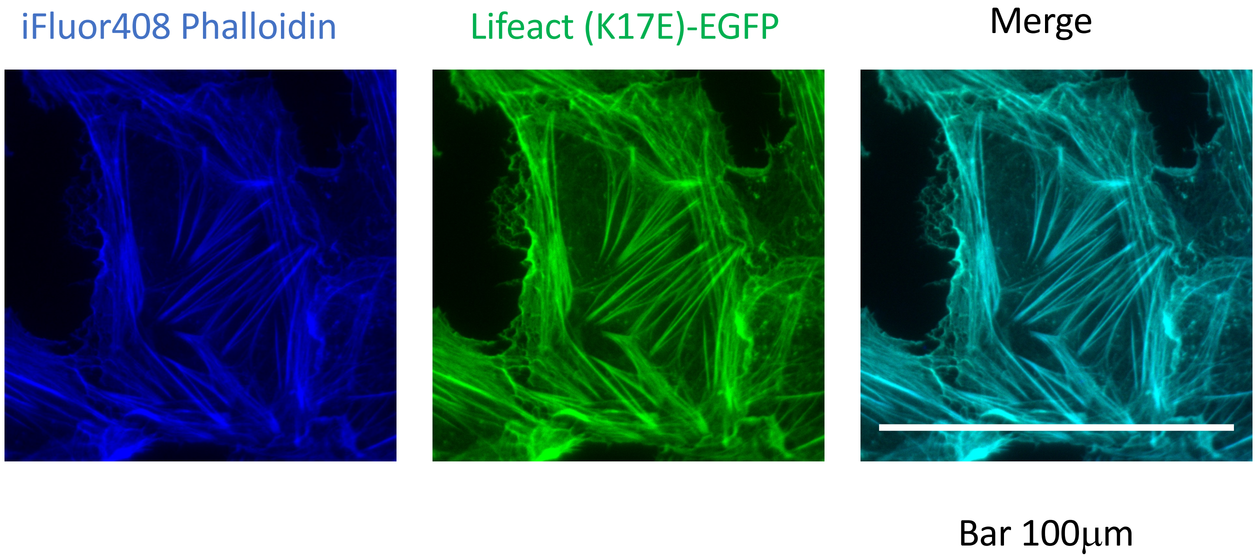


Nagasaki et al., Supp Figure 5

### Supplementary Figure 6

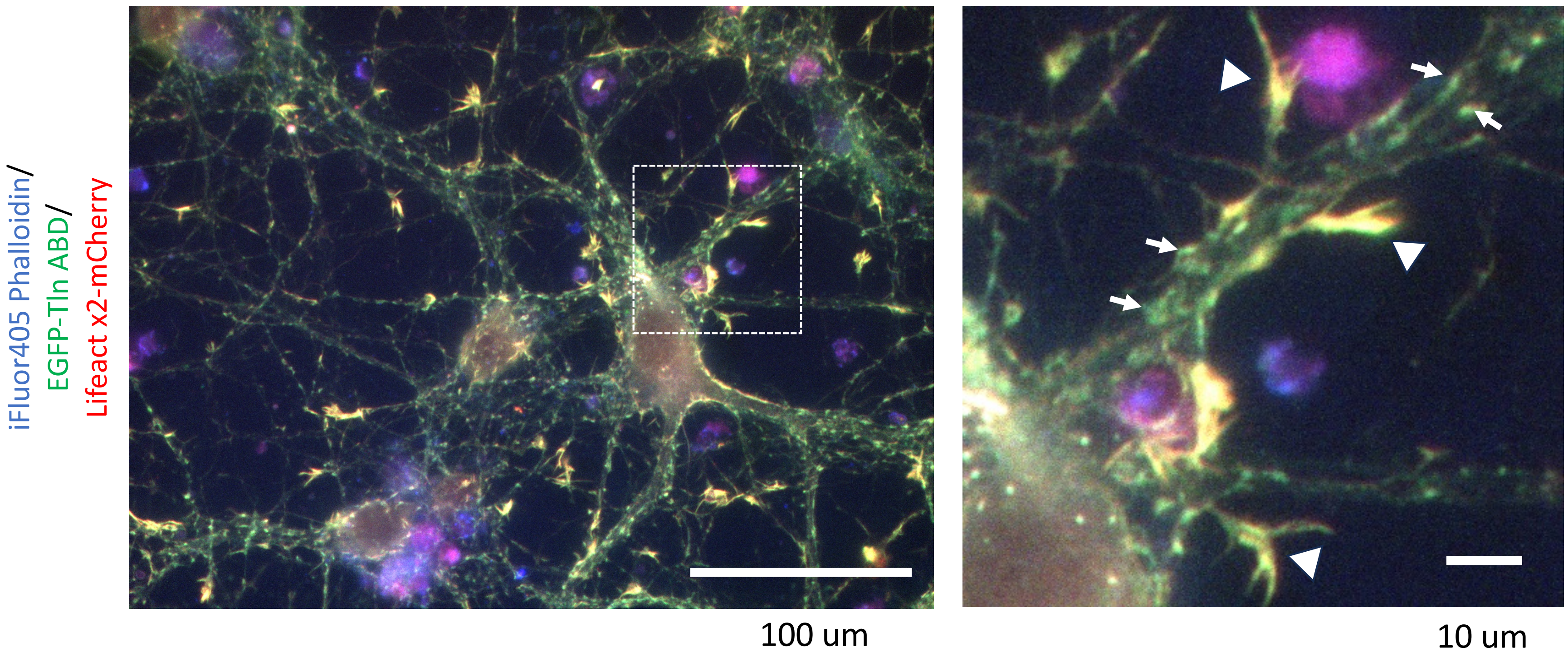


Nagasaki et al., Supp Figure 6

### Supplementary Figure 7

phalloidin-iFluor 405 | Tln ABD -EGFP | mCherry-alpha actinin ABD

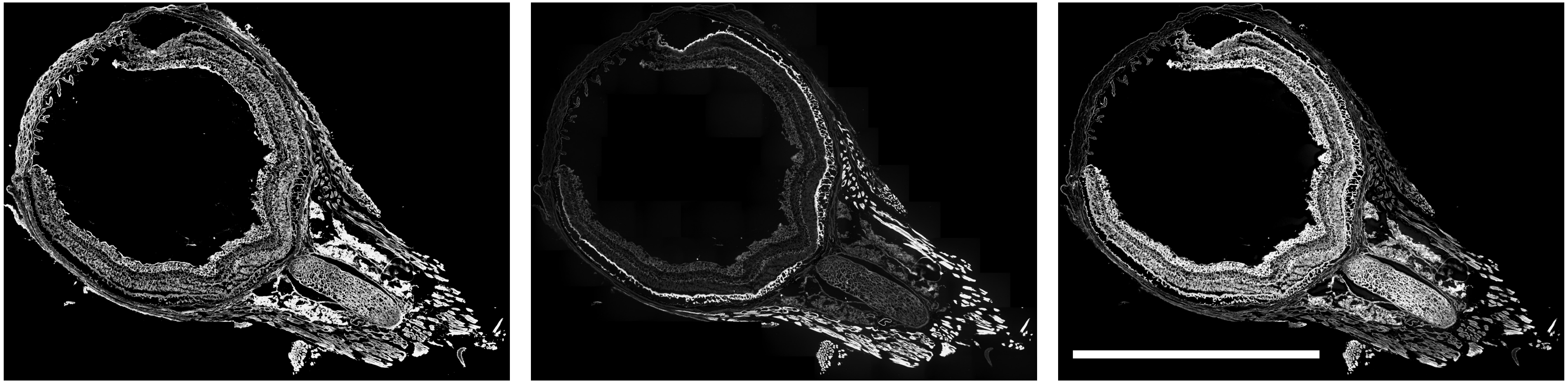

Nagasaki et al., Supp Figure 7

### Supplementary Figure 8

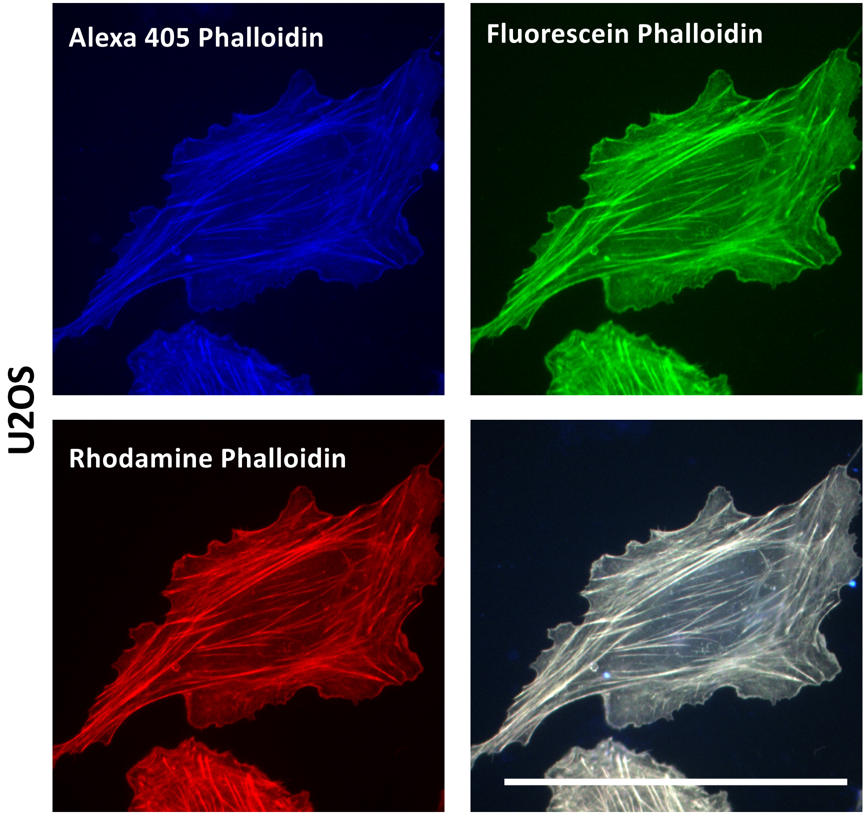


Nagasaki et al., Supp Figure 8

### Supplementary Figure 9

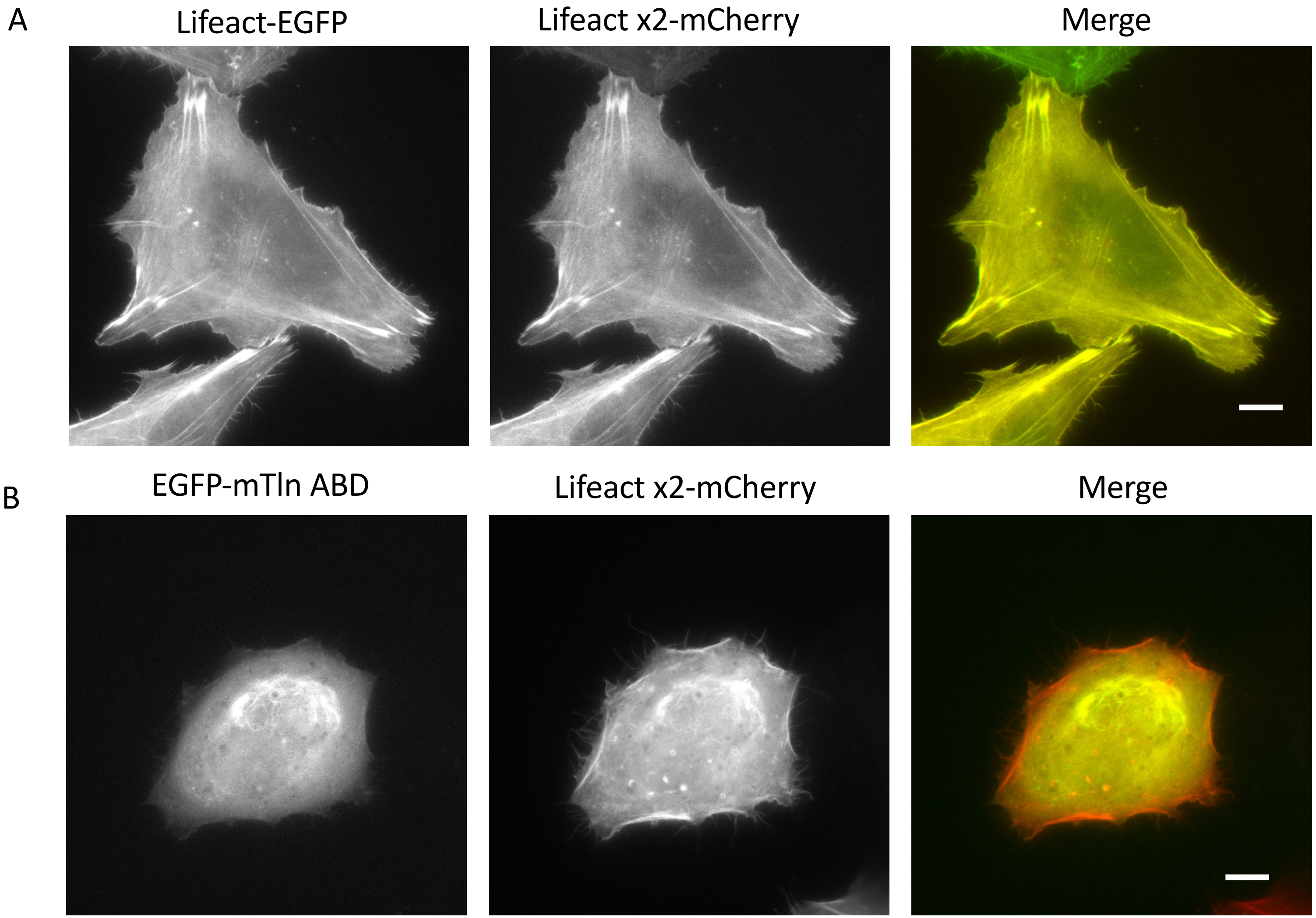


Nagasaki et al., Supp Figure 9
