## Supplementary Table 1 for "Actin Painting: a multicolor fluorescence staining method highlighting cell type-specific differences in the composition of filamentous actin architectures"

Supplementary Table 1. Information on protein-based actin probes.

| | Probe | Species | Sequence | Tags | Reference |
|---|---|---|---|---|---|
| 1 | Lifeact | *S. cerevisiae* | MGVADLIKKAESISKEE | EGFP | Riedl, J. *et al*., (2008) |
| 2 | Lifeact x2 | *S. cerevisiae* | MGVADLIKKAESISKEEGTMGVADLIKKAESISKEE | mCherry | This report |
| 3 | Lifeact (E17K) | *S. cerevisiae* | MGVADLIKKAESISKEK | EGFP | Belyy, A *et al*., (2020) |
| 4 | Rng2 ABD | *S. pombe* | KLDVNVGLSRLQSQAGAPVGTKGSNTRLAAKQRETLQAYDYLCRVDEAKKWI<br>EECLGTDLGPTSTFEQSLRNGVVLALLVQKFQPDKLIKIFYSNELQFRHSDNINK<br>FLDFIHGIGLPEIFHFELTDIYEGKNLPKVIYCIHALSYFLSMQDLAPPLIKSDENLS<br>FTDEDVSIIVRRLRQSNVILPNFKA | EGFP | Hayakawa, Y. *et al*., (2023) |
| 5 | Tln ABD | *M. musculus* | ILEAAKSIAAATSALVKAASAAQRELVAQGKVGAIPANALDDGQWSQGLISAA<br>RMVAAATNNLCEAANAAVQGHASQEKLISSAKQVAASTAQLLVACKVKADQ<br>DSEAMKRLQAAGNAVKRASDNLVKAAQKAAAFEDQENETVVVKEKMVGGI<br>AQIIAAQEEMLRKERELEEARKKLAQIRQQQYKFLPSELRDEH | EGFP | Kost, B. *et al*., (1998) |
| 6 | Cofilin | *D. discoideum* | MSSGIALAPNCVSTFNDLKLGRKYGGIIYRISDDSKEIIVDSTLPAGCSFDEFTKC<br>LPENECRYVVLDYQYKEEGAQKSKICFVAWCPDTANIKKKMMATSSKDSLRKA<br>CVGIQVEIQGTDASEVKDSCFYEKCTKI | mCherry | Umeki , N. *et al*., (2013) |
| 7 | α-actinin ABD | *D. discoideum* | GTPVSGNDKQLLNKAWEITQKKTFTAWCNSHLRKLGSSIEQIDTDFTDGIKLA<br>QLLEVISNDPVFKVNKTPKLRIHNIQNVGLCLKHIESHGVKLVGIGAEELVDKNL<br>KMTLGMIWTIILRFAIQDISIEELSAKEALLLWCQRKTEGYDRVKVGNLHTSFQ<br>DGLAFCALIHKHRPDLINFDSLNKDDKAGNLQLAFDIAEKELDIPKMLDVSDM<br>LDVVRPDERSVMTYVAQYYHHFSASRKAETA | mCherry | This report |
